# Semiautomated detection of *Pseudomonas aeruginosa* from diverse water samples using highly selective Z-broth and a redox potential monitor

**DOI:** 10.1101/2020.02.17.946384

**Authors:** Géza Szita, Szakmár Katalin, Sándor Bernáth, József Sövényi, Gábor Fülöp, Hadjiev Janaki, György Csikó, Norbert Solymosi, László Makrai

## Abstract

*Pseudomonas aeruginosa* is a facultative bacterial pathogen with increasing public health risk potential. Contaminated water in hospital environments is a growing cause of multidrug-resistent *P. aeruginosa* nosocomial infections that are life-threatening and are costly to treat. *P. aeruginosa* is common in natural water bodies, but it is not unusual in drinking water and has been detected in bottled water, also. Suppliers in Europe must eliminate live forms of the bacterium in drinking water to meet human consumption requirements. Laboratory testing for the presence of viable *P. aeruginosa* is mostly done manually using culture media but conductance/impedimetry measurements for detection are also available. In order to improve expedience and cost efficiency as well as to automate the detection and the data registration, we applied the highly selective Z-broth culture media and redox potential monitoring to detect *P. aeruginosa* from water. The Z-broth is based on only a few, stable chemicals that provide consistency of quality and long shelf life. It limits growth to *P. aeruginosa* and, thus, eliminates the need for subsequent microbiological identification steps of the European Standard procedure defined in ISO 16 266:2006 (2018). A redox potential monitor was used in this work, that simultaneously recorded 64 sample curves and automatically marked positive samples reliably within 24 hours after sample initialization. In comparison, the standard method requires additional tests that prolongs identification to some days. Practical applicability and reliability of the method in this paper was demonstrated by testing a total of 739 water samples of which 222 were tap water, 342 well water and 145 had been taken from swimming pools.

It is considered this method is well suited to process large numbers of samples for purposes of detecting *P. aeruginosa* contamination with relative ease, little cost in shortened time. Using Z-broth in combination with redox potential monitoring can be recommended for central laboratories for routine testing of drinking water for live *P. aeruginosa* presence. As well, using this method for testing of fluids and surfaces for *P. aeruginosa* contamination would be advantageous in such environments as the hospital industry, swimming pools and spas, where cleaning routines and *P. aeruginosa* transmission prevention protocols could be made much more resultful and cost-effective due to ease and speed of processing both spot check and water samples.

**Highlights:**

- A method was applied for detection of viable *Pseudomonas aeruginosa* in 739 water samples.
- Redox potential monitoring of Z-broth cultures exclusively registered growth of *P. aeruginosa*.
- The method is highly selective, reliable and partly automated, with lower labor costs.
- The method is suited for testing of water and hospital environments for *P. aeruginosa*.

## 1. Introduction

*Pseudomonas aeruginosa* is an ubiquitous facultative anaerobic Gram-negative bacterium. It is common in our environment in the soil, water, foods, feces, so it can get easily into drinking water.

Testing for the presence of *P. aeruginosa* in drinking water is an important area of interest for this study due to the fact that the European Union’s drinking water standard allows zero *P. aeruginosa* presence (ISO, 2018). Drinking water checks are conducted at a great number of laboratories and vast populations depend on reliability of results.

Beyond the abovementioned significance for drinking water, *P. aeruginosa* is one of only three bacterial species that are listed in the latest WHO global priority pathogens list bulletin. Other than *P. aeruginosa*, only *Acinetobacter baumannii* and certain members of *Enterobacteriaceae* are listed in the critical priority group (WHO 2017). The word „critical” implies that these pathogens are responsible for serious, often fatal septicemia, as well as difficult-to-treat, varied chronic infections. The bulletin has assigned the highest priority of urgency to do research on control of members of the critical priority group (WHO, 2017). *P. aeruginosa* strains are known to complicate pneumonias (Hopkins, 1983), cystic fibrosis (Caskey et al., 2018) and chronic obstructive pulmonary disease (COPD) (Rakhimova et al., 2009) but they are important pathogens in iatrogenic infections, also (Hopkins, 1983. Rakhimova et al., 2009. Zorilla-Vaca 2016). *P. aeruginosa* strains isolated from affected patients may have as low virulence as environmental isolates (Martins et al., 2014) but some isolates from human cases or from the environment were shown to carry numerous virulence-associated genes (Scaccabarozzi et al., 2015).

In the WHO global priority pathogens list bulletin, the need for more research into prevention of *P. aeruginosa* infections gets emphasis as an effective means of reducing morbidity rates and, thus, reducing the spread and scale of antibiotic resistance (WHO 2017).

However, the standard method of detection of viable *P. aeruginosa* is time-consuming because requires culturing on cetrimide-containing selective media and any necessary additional microbiological identification tests. This standard procedure is also labour-intensive; plus it requires personnel having microbiological training background. Furthermore, the cetrimide-agar, with selectivity based on an inhibitory substance may not reliably demonstrate trace contamination with *P. aeruginosa* in every sample (Wesche, 2009).

We, therefore, designed a new method, as described below, that is selective, reliable and mostly automated, as well as it cuts down the time to results of detection, to help organizations involved into provision of drinking water to the public or the health industry. We used a highly selective media, the Z-broth that provides growth support to only *P. aeruginosa* without any inhibitory substance (Szita et al., 2007). Z-broth has been shown to be a suitable means of accelerated selective *P. aeruginosa* detection in a study where bacterial growth was detected by impedance measurement (Erdősi et al., 2012). However, impedance/conductance measurements can be in error due to even slight temperature variations and they also demand high hardware cost and skilled personnel. Redox potential monitoring does not have the drawbacks of impedance measurement. Good performance of the redox potential monitoring method for *Campylobacter* detection has been shown earlier (Erdősi et al., 2018).

## 2. Materials and methods

### 2.1. Redox potential measurement setup

Redox potential was measured by a multichannel reader, a device named ‘MicroTester’ that monitored redox potential in maximum 64 samples simultaneously (4 blocks of 16 tube test units) and registered values using the manufacturer’s Windows-based software (Erdősi et al., 2012). The parameter of decision threshold (DT) was set to be −0.6 mV/min drop rate in this study; this DT has been established previously (Erdősi et al., 2012).

### 2.2. Culture media

Composition of Z-broth is as follows: acetamide (5 g/L), potassium hydrogen phosphate (3 g/L), potassium dihydrogen phosphate (3 g/L), potassium tetrathionate (1 g/L), and magnesium sulphate heptahydrate (0.05 g/L); pH = 7.2. The medium was sterilised by filtration.

As a means to verify presence or absence of bacteria in our examined water samples, we used MPA agar cultures. The composition of MPA agar is: potassium monohydrogenphosphate (7 g/L), potassium dihydrogenphosphate (3 g/L), ammonium sulfate (1 g/L), magnesium sulfate heptahydrate (0.1 g/L), yeast extract (0.1 g/L), palmitic acid (1.5 g/L), Triton X□100 (10 ml g/L) and agar (15 g/L); pH = 7.2. The medium was sterilized by autoclaving at 120°C for 20 min.

### 2.3. Water samples

To check practical applicability of the above testing setup for detecting presence of live *P. aeruginosa* in water, we conducted tests on 739 water samples of which 222 were tap water, 342 well water and 175 samples were collected from swiming pool water. Samples were obtained within the borders of Hungary from various areas. Each sample was taken from different bodies/flows of water. Volumes of 1 L were collected from each source in sterilized glass flasks supplied with 0.5 g sodium thiosulfate to deactivate any oxidizing agent in the water.

Of each 1 L water sample, 100 mL water was filtered through microfilters (Millipore 0.45 μm membrane, diameter 47 mm), and the membranes were transferred into 10 mL Z-broth, each, in the measuring cells of the MicroTester device. Monitoring of redox potential in the measuring cells was conducted for 24 h at 37 °C. For negative control, a sterile Millipore membrane was incubated in 10 mL Z-broth with each run of tests.

### 2.4. Checking for bacteria after test runs

After the incubation of water samples for 24 hours, inoculums were spread on MPA agar plates from each measuring cell. Suspect bacterial colonies were checked for their oxidase reaction, lack of indol production, urease test, glucose oxidation, and reduction of nitrate into nitrogen gas (ISO 2018), to confirm if they were *P. aeruginosa*.

## 3. Results

### 3.1. Monitoring time pre-set for marking absence of P. aeruginosa

The key task in this study was to establish either presence or absence of *P. aeruginosa* trace contamination of water samples. Presence of the bacterium was indicated by redox potential change in the culture fluid. When the redox potential drop rate reached the detection criterion (DC) value in any sample, that sample was marked positive by the registering software automatically. An example of redox potential shift caused by *P. aeruginosa* growth, and the test evaluation principle are illustrated in Figure 1.

**Figure 1.**
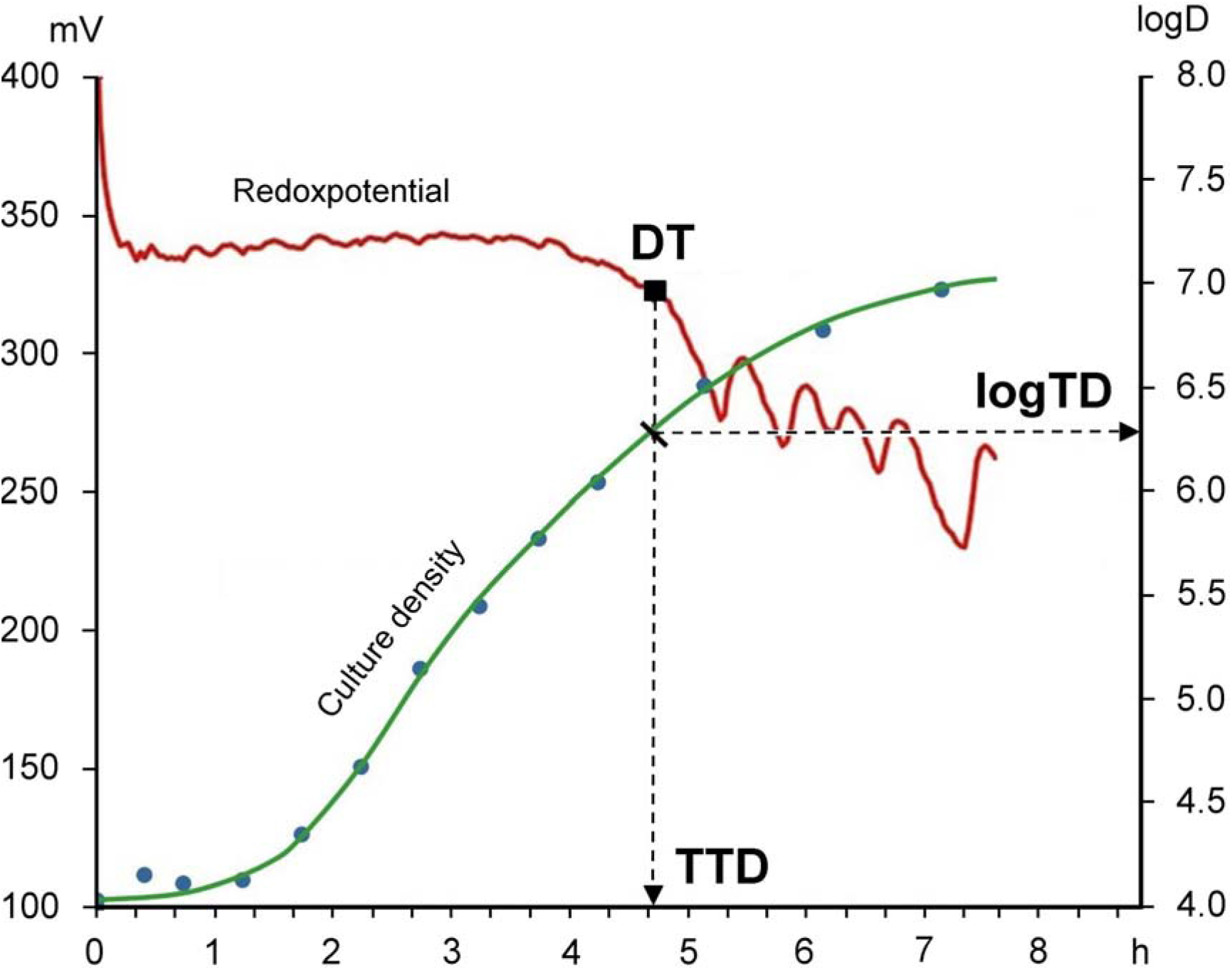
Sample graph of redoxpotential in correlation to culture density. *Pseudomonas aeruginosa* was cultured in Z-broth. The right vertical axis shows the logarithm of culture dendrit in cfu/ml (logD). As logD values increase, the bacterial metabolites start to cause an accelerating drop of redoxpotential. When reaching a certain redoxpotential drop rate, called decision threshold (DT = −0.6mV/min), the test result is deemed to be positive for growth. This DT point gives projections of time to detection (TTD) and the logarithm of threshold density (logTD) the latter of which correlates to the microbe species under the culture conditions. TTD does fluctuate within a certain range with the initial microbe density and only its maximum has assumptive decision value for test result. If the DT is not reached within the maximum TTD that is considered characteristic of the microbe, the test result will be negative. Unchanged redox potential persisting beyond a maximum TTD was taken as proof of absence of the microbe.

In order to find the expectable maximum TTD value characteristic of *P. aeruginosa*, that is, to determine the tipping point to positive/negative readings, a prerun of redox potential monitoring was done. Serial ten-fold dilutions were made starting from a suspension of a *P aeruginosa* colony in physiologic solution. One mL’s taken from each dilution were dispensed into measuring cells of the MicroTester multichannel redox potential monitor, that had been preloaded with 10 mL Z-broth. The measuring cells were incubated in 37 °C water bath while each cell was connected to the monitor. The TTD was 8 h for the sample of undiluted suspension and 10 h, 12 h, 14 h, and 16 h were registered for the sequentially higher dilutions of samples. Bacterial growth could not be demonstrated in samples from higher dilutions. This means that the maximum TTD characteristic of *P. aeruginosa* to give positive result was 16 hours in our system and samples with unchanged redox potential beyond that time limit should have absence of *P. aeruginosa*. These samples could be marked negative. However, a 24 hour culture period was allowed to determine that a sample was negative, in order to preclude any mistake.

### 3.2. Test results of checking the water samples

Test results of growth of *P. aeruginosa* in water samples are presented in Table 1.

**Table 1.** Results of tests for detecting contamination of water samples with live *P. aeruginosa*

| Sample sort | Number of samples | Positive samples when tested by |  |  |
| --- | --- | --- | --- | --- |
|  |  | redox potential in Z-broth | Z-broth culture | MPA agar |
| Tap water | 222 | 2 | 2 | 2 (0.9%) |
| Well water | 342 | 68 | 68 | 68 (20%) |
| Swimming pool water | 175 | 78 | 78 | 78 (45%) |

Complete coincidence of positive or negative results involving a total of 739 samples was seen when live *P. aeruginosa* presence was identified by any of the three detection methods in Table 1. This includes swimming pool water samples which had strong and variable bacterial contamination. This finding confirmed both high selectivity and adequate growth support of Z-broth for *P. aeruginosa*.

## 4. Discussion

Purpose of our work was to develop a fast, semi-automated procedure to detect trace levels of *P. aeruginosa* presence in water. The concept is based on the use of the elective medium, Z-broth and redox potential monitoring.

Both the Z-broth and redox potential monitoring bring their respective particular advantages to the detection procedure.

We have developed Z-broth earlier to be able to selectively grow *P. aeruginosa* without any inhibitory additive in the media. (Szita et al., 1998) It is common to include inhibitory substances, e.g., cetridine, in selective culture medias in order to help *P. aeruginosa* detection. However, these substances decrease the detectability of very low bacterial densities when the bacteria are damaged or have compromised viability (Wesche, 2009). Z-broth, being an ‘elective media’, does not contain any inhibitory substance and allows the division and detection of damaged bacteria that are able to complete division and so improve recovery rate of compromised bacterial cells, and increase test reliability. The only carbon and nitrogen source in Z-broth for microbial growth is acetamide which is the simplest yet chemically stable compound still able to support *P. aeruginosa* growth. The bacterium is able to enzymatically decompose acetamide and utilize it for sythesizing its own substances (Szita et al., 1998).

Selectivity of Z-broth for *P. aeruginosa* was demonstrated in a previous study where numerous bacterial genera have been shown to be unable to grow in Z-broth: *Bacillus, Staphylococcus, Micrococcus, Escherichia, Enterobacter, Klebsiella, Citrobacter, Salmonella, Proteus, Yersinia, Aeromonas, Pleisomonas* (Szita et al., 2007). Even sharper specificity was seen when testing other members of the *Pseudomonas* genus. None other than *P. aeruginosa* grew in Z-broth out of 17 species including *P. putida, P. facilis, P. aceris* (Szita et al., 2007).Similarly, in this study, radically limited nutrient supply in Z-broth appeared to have starved all other components of the common bacterial flora in water samples and, in 739 random water samples, not a single positive test result was seen from other than *P. aeruginosa*.

This work demonstrated that redox potential monitoring has three general advantages over traditional microbial cultures for live *P. aeruginosa* detection: (1) it cuts down the incubation time since negative readings take only the time to detection (TTD) which was about one day here with minimal sample preparation; (2) multiple tests can be run and evaluated simultaneously and automatically; and (3) no additional microbiological identification tests and testing reagents are needed after the monitored culturing process. These result into savings of time and resources.

In comparison with the impedance/conductance measurements, the redox potential measurement process is not error-prone due to slight temperature fluctuations. Furthermore, the redox potential measuring devices cost a fraction of what impedance testers cost. Time requirement of impedance/conductance based testing is similar.

Polymerase chain reaction (PCR) has been used to detect *P. aeruginosa* DNA in tap water and bottled water (Momtaz 2011). Great disadvantages of PCR as compared to methods based on initial culturing are that PCR detects DNA, unconnected to viability of the bacterium, and that the method is costly.

On the basis of the obtained test results, it is concluded that redox potential monitoring of Z-broth cultures is a specific and reliable method for the selective and accelerated detection of live *P. aeruginosa* from water samples. Sensitivity of the method is considered very high: theoretically, it should be able to detect as few as a single viable cell of *P. aeruginosa* per redox potential measuring cell. Additionally, because of the use of the elective Z-broth with no microbial inhibitor additive, the method can detect compromised but viable cells of P. aeruginosa, unlike typical inhibitory media media (Wesche, 2009).

Further advantage of Z-broth is the minimal requirement of very low-cost culture media ingredients that combines with chemical stablility in prepared, sterilized form. Synthetic culture media are gaining use even in research of antimicrobial drug residues due to more reliable research results (Szita et al., 2017).

The apparent advantages of the presented method make it a good choice for the drinking water industries where contamination inside and outside the supply system, and on contaminated surfaces (including perhaps, hands of service workers) must have similar attention in control. Therefore, it is considered that laboratories would benefit from using the above detection method when compared to culturing on cetrimide-containing inhibitory media and on additional complementary microbiological tests. Batchwise processing of samples like, e.g., memebrane filters from 10 water samples can also be done when low/no contamination rate of drinking water samples is expected. Even if some positive batches may need to be re-tested by individual samples, batch tests would raise lab productivity greatly. Such batchwise processing is impossible when using culture plates as per the ISO standard (ISO, 2018).

Research conducted in health industry establishments demonstrated that multidrug and carbopenem resistant *P. aeruginosa* strains are increasingly frequent in nosocomial infections, especially in the intensive care units (Obritsch et al., 2005, Barna 2014). Few data are revealed publicly on costs of treatments of nosocomial *P. aeruginosa* infection, costs of infection control, or about the liability burdens from chronic or fatal *P. aeruginosa* nosocomial infections but they can be estimated to be significant (Nathwani et al., 2014). Investigations into nosocomial cases brought attention to high rate of *P. aeruginosa* transmission by contaminated water (Breathnach et al., 2012, Salm et al., 2016, Amoreux et al., 2017, Devarajan 2017). Washing sinks, showers, toilets used by patients were often contaminated with *P. aeruginosa* even if cleaned twice daily using quternary-ammonium based formulations (Salm et al., 2016, Amoreux et al., 2017). Thus, conducting regular spotchecks would be essential in disease prevention protocols to help reduce transfer of dangerous nosocomial pathogenic *P. aeruginosa* strains and to reduce associated morbidity and mortality rates as well as help slow down the spread of multipledrug-resistant *P. aeruginosa* strains in the health industry. Infectious agent transmission is a problem also in swimming pools and public baths where humidity can support survival and biofilm formation of *P. aeruginosa* (Rice 2012 et al., Proctor 2018 et al., Tirodimos 2018 et al.,). The method described in this paper can greatly decrease costs and time to identify surfaces contaminated with *P. aeruginosa* and to verify desinfection. In our work, water samples were tested. However, when the samples are e.g., swabs of sink surface, of shower walls, drain openings and alike, the sample swabs can be simply dropped into Z-broth in the redox potential measuring cells and automatically processed to determine viable *P. aeruginosa* presence with the same good work and time efficiency as water samples. Swabs can be processed batchwise, also.

## 5. Conclusions

- The results allow the conculsion that the procedure described in this study is a simplified, specific, reliable and relatively inexpensive method for detecting viable *Pseudomonas aeruginosa*.
- The method is suited for testing large numbers of varied water samples or swabs.
- The method can be adapted as a feasible routine component of preventive hygiene and desinfection protocols to reduce levels of *P. aeruginosa* contaminations and infections.

## Acknowledgement

This research was funded using regular budget resources of the Department of Food Hygiene, University of Veterinary Medicine, Budapest, Hungary.

## Abbreviations

DT: decision threshold
TTD: time to detection
WHO: World Health Organization

